# Silica particles trigger the production of exopolysaccharides in harsh environment plant growth-promoting rhizobacteria and increase their ability to enhance drought tolerance

**DOI:** 10.1101/2020.05.20.106948

**Authors:** Anastasiia Fetsiukh, Julian Conrad, Jonas Bergquist, Fantaye Ayele, Salme Timmusk

**Affiliations:** Department of Forest Mycology and Plant Pathology, Swedish University of Agricultural Sciences, Uppsala, Science for Life Laboratory, Sweden; Swedish National Cryo-EM Facility, Science for Life Laboratory, Sweden; Department of Analytical Chemistry, Uppsala University, Sweden

## Abstract

In coming decades drought is expected to expand globally owing to increased evaporation and reduced rainfall. In order to reduce the vulnerability of agricultural systems we need to understand the crop plant growth environment. Understanding, predicting and controlling the rhizosphere has potential to harness plant microbe interactions, improve plant responses to environmental stress and mitigate effects of climate change. Our plant growth-promoting rhizobacteria (PGPR) are isolated from the natural laboratory ‘Evolution Canyon’ Israel (EC). The endophytic rhizobacteria from the wild progenitors of cereals have been co-habituated with their hosts for long periods of time. The study revealed that silica particles (SN) triggered the PGPR production of exopolysaccharides (EPS) containing D-glucuronate (D-GA). This leads to increased plant biomass accumulation in drought-stressed growth environments. The PGPR increased EPS content increases the water holding capacity (WHC) and osmotic pressure of the biofilm matrix. Light- and electron-microscopic studies show that in the presence of SN particles, bacterial morphology is changed, indicating that SNs are associated with significant reprogramming in bacteria.

The results here show that the production of EPS containing D-GA is induced by SN treatment. The findings encourage formulation of cells considering microencapsulation with materials that ensure higher WHC and hyperosmolarity under field conditions. Our results illustrate the importance of considering natural soil nanoparticles in the application of PGPR. Osmotic pressure involvement of holobiont cohabitation is discussed.

## INTRODUCTION

Agriculture faces several challenges at the global level and in coming decades, drought is expected to expand globally owing to increased evaporation and reduced rainfall or changes in the spatial and temporal distribution of rainfall(Dai 2012). The scientific community across the world is earnestly looking for novel solutions for enhancing crop plant stress tolerance under limited resource availability, and several environmentally friendly solutions have shown huge potential but need to optimised for wide scale field application. One such solution includes strengthening plant natural defence systems by soil microbes. In order to reduce agriculture vulnerability to climate change we need to understand the crop plant rhizosphere, which is one of the most complex microbial consortia on earth. Predicting and controlling the rhizosphere has a potential to harness plant microbe interactions and restore plant ecosystem productivity, improve plant responses to environmental stress and mitigate effects of climate change (Xu, Naylor et al. 2018, Fitzpatrick, Mustafa et al. 2019).The first report on PGPR-induced drought stress was published in 1999 (Timmusk and Wagner 1999) and its reported change in ERD15 and RAB18 gene expression has been repeatedly discussed (Timmusk and Wagner 1999, Sukweenadhi and al 2015, Naylor and Coleman-Derr 2017, Bandeppa, Paul et al. 2019, Shimaila and Glick 2019, Timmusk, Copolovici et al. 2019). It is known that the rapid nature of changes in climatic pattern does not allow the adaptive and supportive crop plant microflora development but rather the plant microbiome, which is evolved with its host and can significantly contribute to its host stress adaptation (Liepe, Hamann et al. 2016, Fitzpatrick, Copeland et al. 2018, Fitzpatrick, Mustafa et al. 2019). Our wild barley *(Hordeum spontaneum)* endophytic isolate *P. polymyxa A* 26 originates from a habitat exposed to various stress factors: the Evolution Canyon (EC) South Facing Slope (SFS)(Timmusk, Paalme et al. 2011, Timmusk, Abd El-Daim et al. 2014) (Fig 1). The *P*. *polymyxa* A26 Sfp-type 4-phosphopantetheinyl transferase deletion mutant strain (A26Sfp), lacking the ability to produce both nonribosomal peptides and polyketides, was created and we showed that the Sfp-type PPTase gene in *P*. *polymyxa* is a gate-keeper for the bacterial drought tolerance enhancement (Timmusk, Kim et al. 2015). EC is a well described natural laboratory where microbes have co-habituated with the hosts over a long period of time (Nevo 2012). Reciprocity between the organisms is generally accepted (Gilbert, Bosch et al. 2015, Gilbert 2016, Rice, Kallonen et al. 2016, Gilbert 2019) and organisms that have co-evolved within the environment are more robust to environmental stress situations (Robinson, Klein et al. 2018, Vega 2019). Microbes found in extreme habitats developed a different strategy to adapt to such conditions through evolution. Holobiotic complex relationships in the given framework, not robotic associations of genes, are becoming more and more accepted in developmental and evolutionary biology (Gilbert, Bosch et al. 2015, Gilbert 2016, Rice, Kallonen et al. 2016, Gilbert 2019). Hence, understanding the framework of adaptation to extreme environments is gaining interest in a period of environment change (Robinson, Klein et al. 2018, Vega 2019).

**Figure 1.**
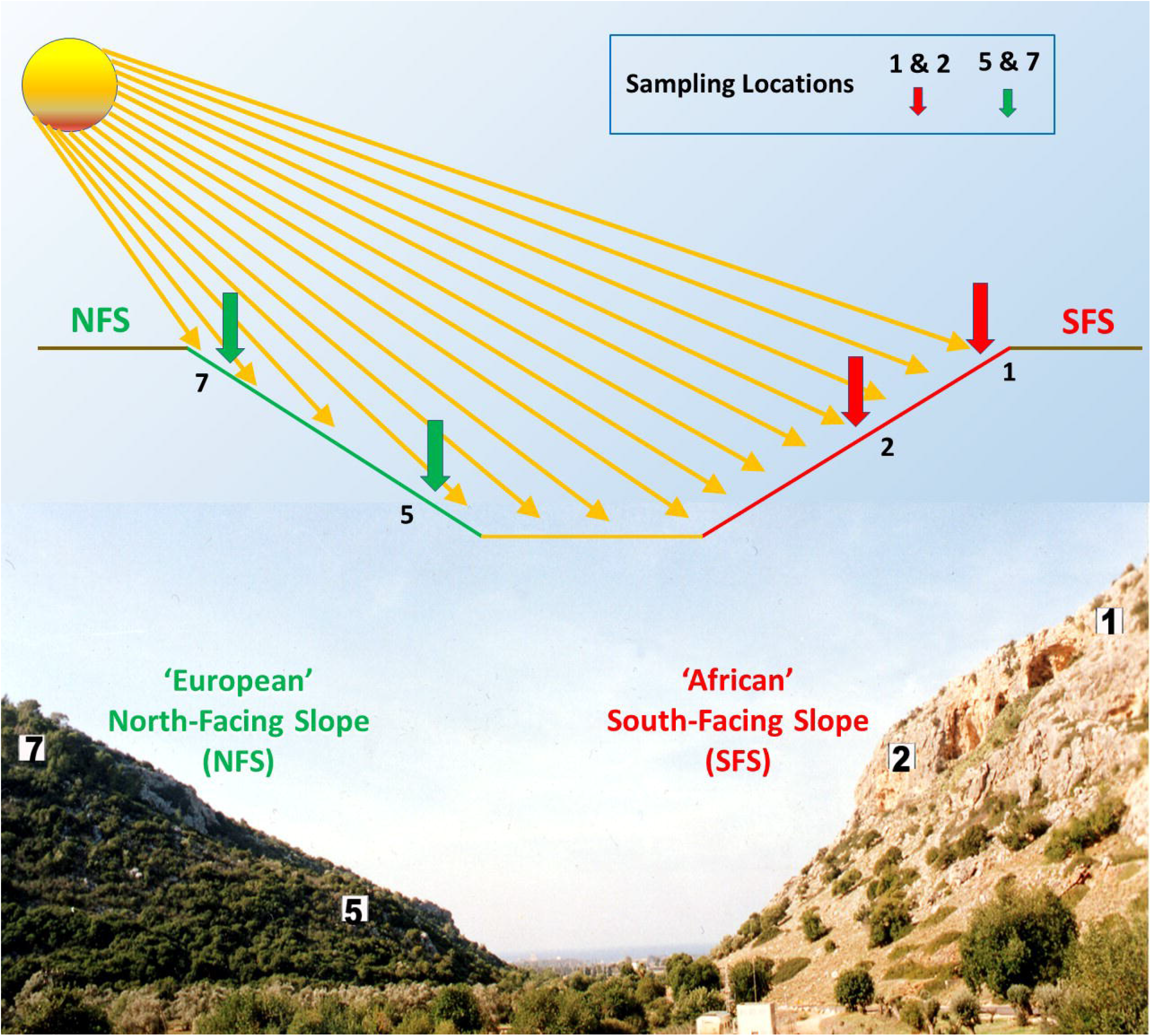
The Evolution Canyon (EC) model. Schematic diagram of the Evolution Canyon at Lower Nahal Oren, Mount Carmel (source^10^: Nevo, 2012 Evolution Canyon,” a potential microscale monitor of global warming across life, PNAS 109; 8) (Photo by S. Timmusk).

Our recent plant nanointerface interaction research focusing on native soil particles shows that silica nanoparticles (SN) promote PGPR attachment and plant colonization, and this effect is translated into crop yield enhancement under stress conditions (Timmusk, Seisenbaeva et al. 2018). The solid phase of any soil is composed of minerals (inorganic) and organic material. Minerals predominate in virtually all soils. The most abundant class of minerals are silicates, which have a substantial impact on soil characteristics, as their surfaces are inherently reactive, potentially forming strong or weak chemical bonds with soluble substances and further regulating the composition of the soil solution (Luyckx, Hausman et al. 2017). Silica particles have been studied as effective agents for phytoremediation, pathogen resistance and as silicon fertilizer (Karunakaran, Suriyaprabha et al. 2013, Rangaraj, Gopalu et al. 2014, Luyckx, Hausman et al. 2017). The mechanism of their action still needs further studies prior thier efficient wide scale application Several forms of commercial SNs have been produced and well characterised (Various 2006, Orts-Gil, Natte et al. 2011). The interaction between plant and bacterial inoculants is coordinated by the complex reactions on the surface of root. The bacterial exopolysaccharide (EPS) matrix in the immediate vicinity of cells serves as the interface with the surrounding environment and root.

Here we study the effect of commercial silica particles on A26 and A26Sfp in the culture medium, hydroponic- and soil system. We show that SNs trigger EPS production which increases the water holding capacity and osmotic pressure of the bacterial biofilm. Our light and electron microscopic studies show that SN effect is associated with bacterial cell elongation and aggregate formation.

## RESULTS

### EPS production and EPS D-GA content with and without SNs in bacterial growth medium

Strains A26, A26Sfp were grown in 1/2 TSB with 50 μg ml^-1^ silica particles (SN) at 30 ±2 °C for 24 hours. While the SNs didn’t have significant impact on the bacterial number, the nanoparticles improved A26 and A26Sfp EPS production by 46 and 29 % respectively (Table 2). A26Sfp EPS production was 30-40 % higher than its wild type and A26Sfp SN treatment caused a further 20% increase in EPS production (Table 2).

**Table 1.** Summary of plant treatment assays. Randomized block design was applied to winter wheat *(Triticum aestivum* cv. Stava) PGPR treatments in peat (Sol mull, Hasselfors), and hydroponic culture. Data were subjected to statistical analysis as described in ‘Data confirmation and validation’

| Time (days) | 1 | 5-8 | 8 | 8-15 | 15 |
| --- | --- | --- | --- | --- | --- |
| Seed sterilization | Ethanol, NaOCl and sterile water |  |  |  |  |
| Peat soil system |  | Inoculation A26/A26SN<br>A26Sfp<br>A26SfpSN |  |  |  |
|  |  |  | Rhizobacterial population assay |  |  |
|  |  |  |  | Drought treatment |  |
|  |  |  |  |  | Harvest/Plant biomass analysis |
| Hydroponic system |  | Inoculation A26/A26SN<br>A26Sfp/<br>A26SfpSN |  |  |  |
|  |  |  | Bacterial population assay/<br>EPS, D-GA assay |  |  |
| Bacterial cell culture | A26/A26SN<br>A26Sfp/<br>A26SfpSN cultures | EPS, D-GA assay |  |  |  |

**Table 2.** EPS and D-GA assessment in relation to bacterial growth

|  | Bacterial<br>population<br>log CFU/ml<sup>1</sup> | EPS titre<br>(µg/ml) | EPS D-GA<br>content<br>(%) | D-GA (E-03<br>µg/ml) |
| --- | --- | --- | --- | --- |
| ½ TSB<br>cultures |  |  |  |  |
| A26 | 9.00±0.4<sup>a2</sup> | 11±2<sup>a</sup> | 3 | 0.3±0.03<sup>a</sup> |
| A26Sfp | 8.69±0.37<sup>a</sup> | 14±2.1<sup>b</sup> | 6 | 0.4±0.04<sup>b</sup> |
| A26SN | 9.0±0.4<sup>a</sup> | 14.5±2.3<sup>b</sup> | 5.5 | 0.8±0.07<sup>c</sup> |
| A26SfpSN | 8.89±0.4<sup>a</sup> | 18±2.4<sup>c</sup> | 5.4 | 0.9±0.08<sup>d</sup> |
| Hydroponic<br>culture |  |  |  |  |
| A26 | 3.15±0.15<sup>b</sup> | 10±1.5<sup>a</sup> | 3 | 0.3±0.07<sup>a</sup> |
| A26SN | 3.09±0.17<sup>b</sup> | 14.8±1.1<sup>b</sup> | 5.8 | 0.5±0.08<sup>b</sup> |
| A26Sfp | 3.08±0.13<sup>b</sup> | 14.6±1.3<sup>b</sup> | 5.8 | 0.8±0.16<sup>c</sup> |
| A26SfpSN | 3.02±0.15<sup>b</sup> | 18±1.9<sup>c</sup> | 5.6 | 1.17±0.2<sup>d</sup> |
| Root wash<br>Control | 1.19±0.11<sup>a</sup> | 0.08 ±0.16<sup>d</sup> | 6 | 0.05±0.01<sup>e</sup> |
| Peat soil |  |  |  |  |
| A26 | 3.02±0.15<sup>b</sup> | ND | ND | ND |
| A26SN | 3.07±0.13<sup>b</sup> | ND | ND | ND |
| A26Sfp | 3.09±0.17<sup>b</sup> | ND | ND | ND |
| A26SfpSN | 3.11±0.13<sup>b</sup> | ND | ND | ND |

<sup>1</sup> The bacteria were re-isolated by selective plating and confirmed by PCR (Timmusk, El Daim et al. 2014). <sup>2</sup>Different letters indicate statistically significant differences (p<0.05). See Material and Methods.

The A26 mutant D-GA content was 30% higher than its wild type. The SNs improved the D-GA content of A26, A26Sfp by 2.7and 2.4 times, respectively. D-GA comprised 5.6-5.8 % of A26SN, A26SfpSN of EPS while the content was 3% in A26 without SNs (Table 2). The D-GA content in A26Sfp was significantly higher than in A26. i.e. 5.8 % of A26 EPS (Table 2).

### EPS production and EPS D-GA content with and without SNs in plant root assay

In order to follow the A26 and its mutant A26Sfp EPS and its D-GA content under more natural settings, the roots of the eight-day-old seedlings were carefully removed from the hydroponic solution. We aimed to collect the biofilms developed after the PGPR inoculation.

The biofilm bacteria were re-isolated by selective plating, confirmed by PCR as described earlier (Timmusk, Abd El-Daim et al. 2014), quantified, and the supernatant polysaccharides isolated and D-glucuronate content recorded (Table 2). A26Sfp and its wild type A26 (10^3^ per ml) were re-isolated in the root biofilms (Table 2).

Significantly higher EPS contents were detected in A26SN and A26SfpSN biofilms in comparison to A26 and A26Sfp biofilms (38 and 13% respectively) (Table 2). While neither the A26 population density nor that of A26SN varied significantly, the D-GA content of the A26SN) was about 40% higher. The D-GA comprised 5.4-5.8 % of A26SN and A26SfpSN while the A26 D-GA content was 3% without SN treatment, and the A26Sfp D-GA content was 6% (Table 2). Additionally, the EPS titre and its D-GA content in the native biofilm layers of the control plant root were evaluated (Table 2). The native biofilms of all the controls comprised around 6% D-GA (Table 2).

### MALDI mass spectrometry

We were interested to learn if the relative increase in EPS titre both in the bacterial cultures as well as in root biofilm assays (Table 2) is reflected in the EPS mass spectrometric analysis. The bacterial cultures were grown in half strength TSB until the logarithmic growth phase and polysaccharides isolated. Typical pattern of A26Sfp and A26SfpSN abundance is shown on Fig 2. Comparative analysis of mass spectra indicates similar pattern of oligosaccharide chains (Fig 2). Quantification shows that while A26SfpSN oligosaccharides with lower m/z values are decreased the oligosaccharides with higher m/z values are increased up to 40% reflecting the change in EPS titre (Table 3).

**Figure 2.**
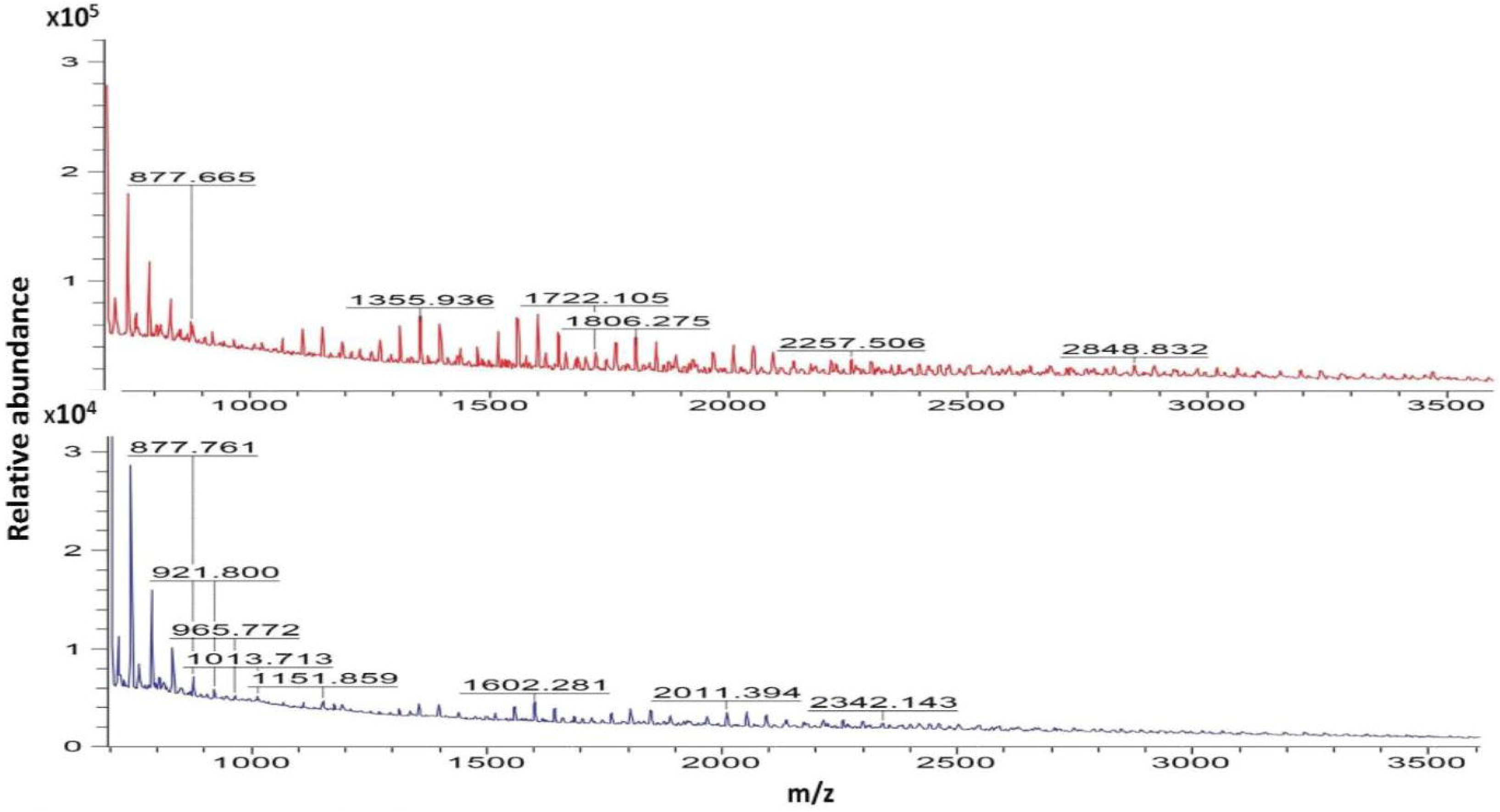
MALDI-TOF mass spectrometry of EPS extracts. Spectra of SN induced A26Sfp extracts (A) compared to A26Sfp EPS extracts (B). Asterisks indicate oligosaccharide chain relative abundance compared in detail in Table (MALDI).

**Table 3.** *Paenibacillus polymyxa* A26Sfp and A26Sfp SN culture filtrate^1^ polysaccharide MALDI-TOF analysis

| № | m/z | Intensity |  | Percentage of change |
| --- | --- | --- | --- | --- |
|  |  | A26Sfp | A26SfpSN |  |
| 1. | 789 | 141907.458 | 90814.536 | 55 |
| 2. | 833 | 88808.125 | 63286.470 | 40 |
| 3. | 877 | 72049.000 | 63445.000 | 14 |
| 4. | 1109 | 45547.000 | 56170.000 | 19↑ |
| 5. | 1151 | 46583.000 | 51814.284 | 10↑ |
| 6. | 1194 | 43345.000 | 44568.000 | 3↑ |
| 7. | 1272 | 36336.000 | 46047.000 | 21↑ |
| 8. | 1356 | 44012.000 | 60857.115 | 28↑ |
| 9. | 1398 | 41128.571 | 53392.158 | 23↑ |
| 10. | 1440 | 35220.000 | 38714.000 | 9↑ |
| 11. | 1476 | 30765.000 | 40044.000 | 23↑ |
| 12. | 1518 | 34532.000 | 48457.729 | 29↑ |
| 13. | 1560 | 39376.080 | 59106.113 | 39↑ |
| 14. | 1602 | 41044.232 | 59972.852 | 32↑ |
| 15. | 1644 | 37606.941 | 48760.800 | 23↑ |
| 16. | 1806 | 32269.259 | 38433.907 | 16↑ |
| 17. | 1848 | 30604.477 | 33294.678 | 8↑ |
| 18. | 2052 | 27555.650 | 32217.000 | 14↑ |
| 19. | 2173 | 25381.000 | 24539.000 | 5↑ |
| 20. | 2215 | 26874.000 | 28241.000 | 5↑ |

<sup>1</sup>See Material and Methods

### Silica particles induce bacterial elongation and aggregate formation

Based on the results described above it was hypothesised that the presence of SN change the bacterial strain development and performance. The hypothesis was studied in the culture media using a Celestron PentaView light microscope with LCD screen and by Talos Arctica cryoelectron microscopy with Falcon III camera. The A26 and A26Sfp cultures with and without SNs were examined from the mid-logarithmic until late-logarithmic phases of growth.

While 40-50 % of SN treated A26 and A26Sfp cells formed elongated agglomerates about 5% elongated cells were formed without SN (Fig 3). Around the elongated cells the bacterial clumps were formed (Fig 3 and Video S1). Vigorous bacterial movements were recorded during the entire logarithmic phase of growth of SN-amended bacteria, while the bacterial movement in the media without SNs was significantly slower after reaching the mid-logarithmic growth (Video S1).

**Figure 3.**
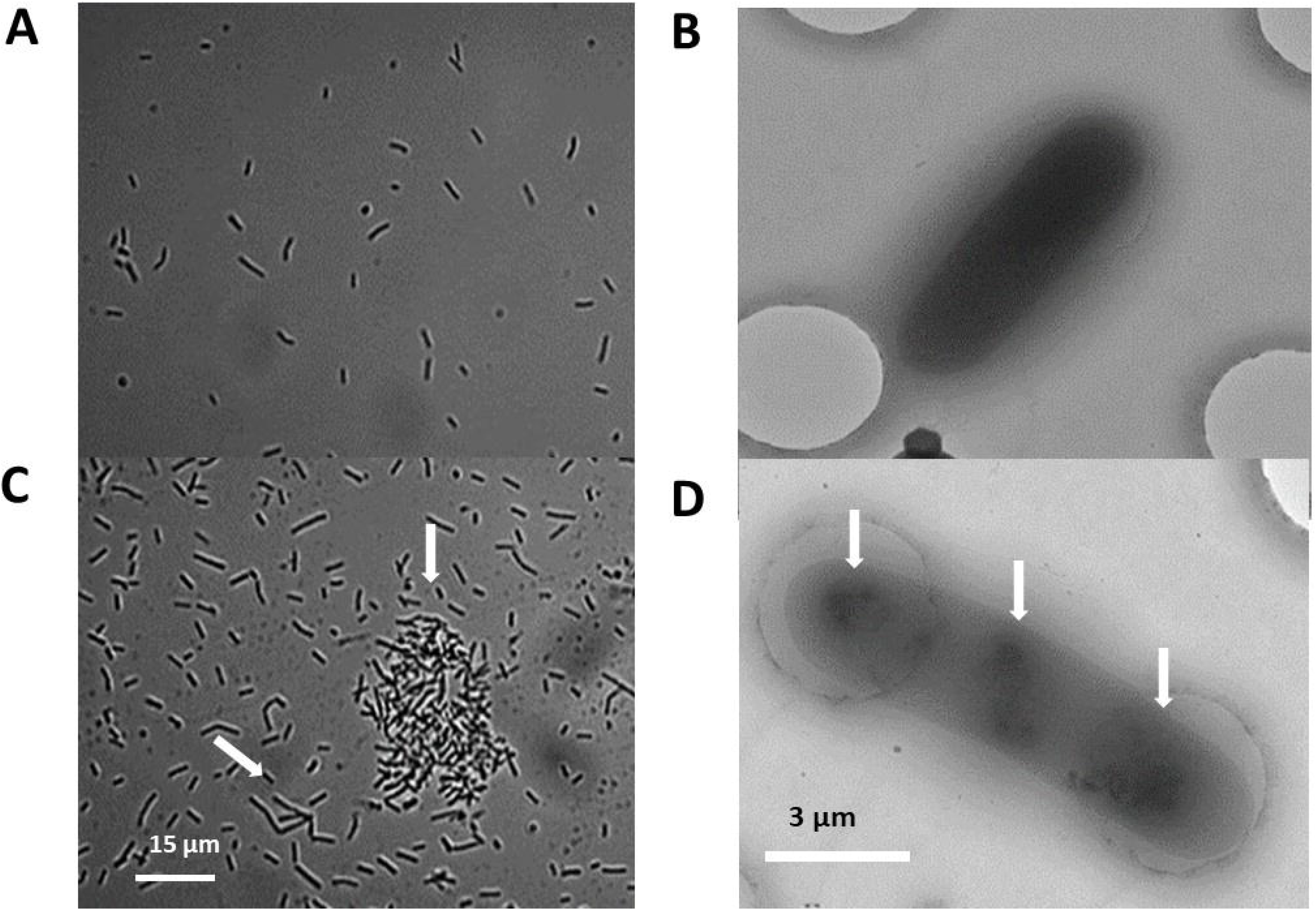
Micrographs of mid-logarithmic bacterial cells grown with and without silica nanoparticles. (SN). Typical light and cryo-electron microscopic images of A26Sfp cells (A, B) and A26SfpSN cells (C, D). Arrows indicate cell elongation, bacterial aggregate formation and changes in nucleoid structure. See Material and Methods.

Bacterial clumps were trapped on TED PELLA individual wells. No clumps but only individual cells were observed without SNs during the logarithmic phase of growth. Cryo-microscopic images of the washed and sonicated clumped cells show longer and larger cells with clear nucleoid structures (Fig 3).

### EPS WHC and osmotic properties

A26 and its mutant A26Sfp, with and without SNs, were grown and EPS extracted as described above. Addition of 0.6% by volume of the biopolymers significantly increased the WHC of peat soil. The SN treatment increased the WHC excretion from A26 and A26Sfp strains by 14 and 16%, respectively (Table 4).

**Table 4.** Effect of bacterially produced biopolymers on osmotic properties^1^ and WHC^1^. Different letters indicate statistically significant differences (p<0.05)

| Strain | Osmolarity mOsm/kg | WHC% |
| --- | --- | --- |
| A26 | 285±13 <sup>a</sup> | 45±3 <sup>a</sup> |
| A26Sfp | 295±10 <sup>a</sup> | 48±3 <sup>a</sup> |
| A26SN | 325±15 <sup>b</sup> | 59±3 <sup>b</sup> |
| A26SfpSN | 347±13 <sup>b</sup> | 71±3 <sup>b</sup> |

<sup>1</sup> See Material and Methods

A26Sfp and its wild type A26 osmotic properties with and without SN treatment were evaluated in 1/2TSB medium grown until mid-logarithmic phase. The contribution of the bacterial polymer to the osmotic pressure was calculated by subtracting the measured osmotic pressure of the pure medium. SN treatment increased the bacterial isolate polymer osmolarity about 15 % (Table 4) and the effect was correlated with EPS production and plant biomass accumulation (r > 0.88 p < 0.01).

### Enhancement of seedling drought tolerance by PGPR with and without SNs

In order to study how SN presence in the PGPR growth medium influences plant growth under drought stress, eight-day-old seedlings were exposed to a seven-day drought stress, harvested and dried. In contrast to the slight but not significant improvement under unstressed condition, 45% biomass improvement was observed by A26SfpSN and 40% improvement by A26SN inoculation under drought stress (*p* <0.01) (Fig 4). Hence, the bacterial strain SN treatments resulted in significant plant shoot dry weight increases (Fig 1). Two to three times increase in the biomass of wheat seedlings treated with A26 and A26Sfp without SNs was observed (Fig 1). The fate of the A26 and A26Sfp with and without SNs was followed using selection plates and by PCR. 10^3^-10^4^ bacterial cfu per gram of plant material after eight days of plant growth was detected (Table 2). When SN formulation was used, statistically similar colonisation number 10^3^-10^4^cfu were recorded (Table 2). Regression analysis indicates that there is a positive interaction between SN-formulated PGPR drought tolerance enhancement activity and EPS D-GA content (r > 0.82 p < 0.01).

**Figure 4.**
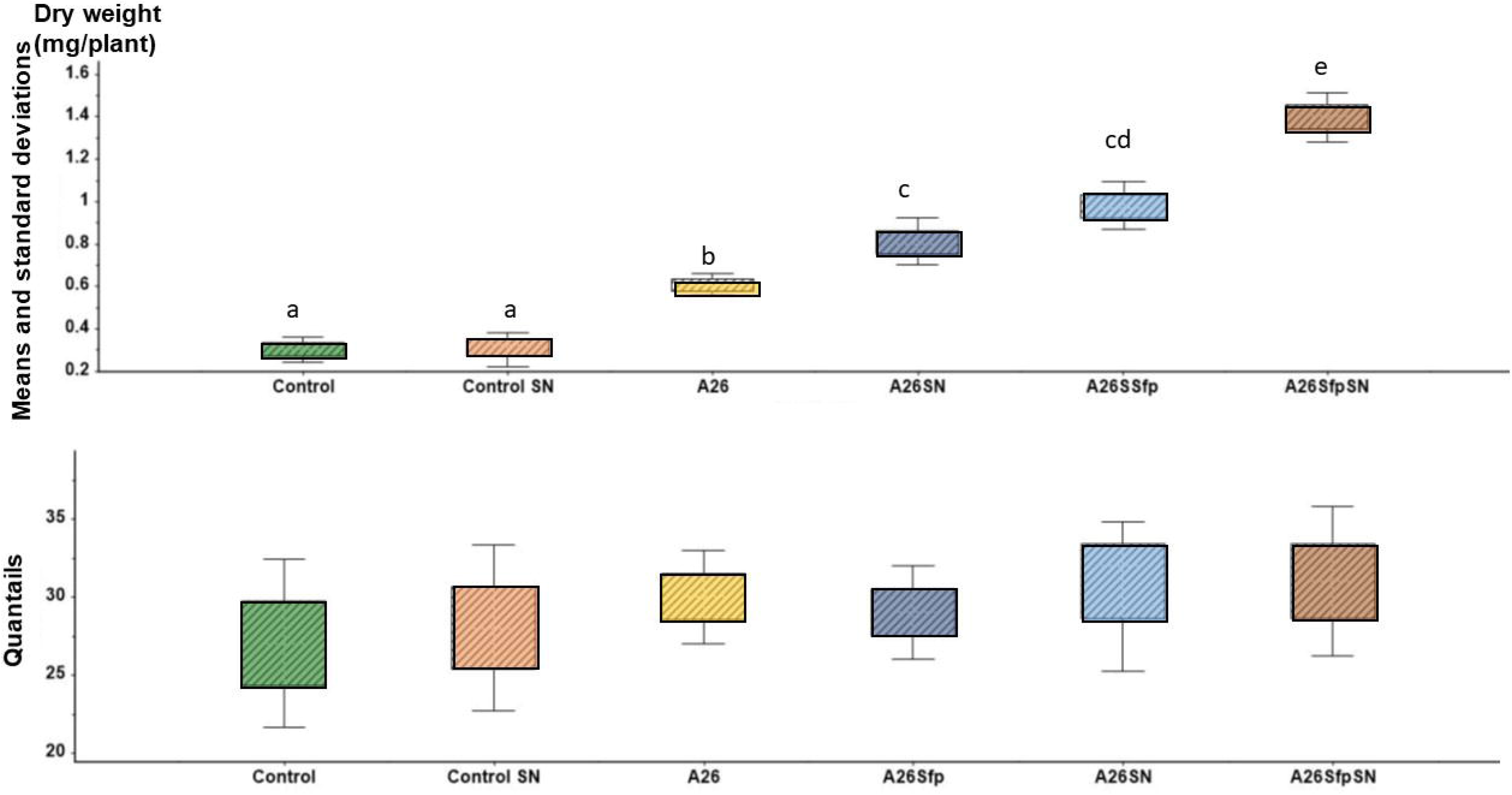
Seedlings weight analysis in soil assay. Plant dry weight means and standard deviations along box plot skewness of A26 A26SN A26Sfp and A26SfpSN treated seedlings. ANOVA univariate analysis was performed and post-hoc LSD tests used to control (p < 0.05). Different letters indicate statistically significant differences.

## DISCUSSION

Here we evaluated the effect of commercial silica nanoparticles (SN) on the EPS production by the harsh environment isolate A26 and its mutant A26Sfp. Although nanotechnology has revolutionized the world, there are safety measures, which should be taken into consideration. Due to the small size, they have properties, which can cause severe damage to ecosystems and human heath (Hristozov, Pizzol et al. 2018). Here we apply silica particles with an idea that SNs are always abundantly present in agricultural systems and therefore present a class of NPs with significantly higher potential for sustainable field applications than metallic NPs. While synthetic NPs are irreplaceable in controlled environment studies, technologies using natural Si nanoparticles are likely to present a possibility wide scale field application.

Earlier we have shown that the stressful SFS slope contains significantly higher populations of bacteria that contain ACCd, form biofilms, solubilize phosphorus, and are tolerant to osmotic stress (Timmusk, Paalme et al. 2011)(Fig 1). The deletion in the Sfp type 4-phosphosphopantheinyl transferase gene resulted in about two-fold enhanced EPS production, the effect reported to correlate with the drought tolerance enhancement of the mutant, and the gene was considered as a key player in the abiotic stress tolerance enhancement (Timmusk, Kim et al. 2015). It has been shown earlier that the strains used in the study affect plant fitness under various stress situations (Timmusk, Abd El-Daim et al. 2014, Timmusk, Kim et al. 2015, Timmusk, Seisenbaeva et al. 2018). Experiments in the study were performed in the bacterial growth medium, hydroponic system and in peat soil (Table 1 and 2). The SNs significantly improved A26 and A26Sfp EPS production in both framework systems and the EPS production in the bacterial growth medium and the root assay (Table 2)

It is well known that even though bacteria are crucial for any biofilm formation, their quantification is insufficient to quantify biofilms. Yet also, owing to the huge complexity of EPS components, their detailed quantification in biofilms is a challenge. We have previously applied D-glucuronate (D-GA) as a proxy for biofilm comprehensive screening, as uronic acid is widely determined as representative of myco-polysaccharides in biofilms (Mojica, Elsey et al. 2007, Timmusk, Seisenbaeva et al. 2018, Timmusk, Copolovici et al. 2019). Here we show that significant differences were detected in A26 and A26Sfp D-GA content dependent on the presence of SNs in the growth medium (Table 2). The D-GA comprised about 6% of EPS in A26 and A26Sfp in comparison to 3% in strains cultured without SNs (Table 2). A26Sfp without SNs comprises 6% of the D-GAs. The higher EPS D-GA content correlates well with the ability of the strain to induce plant drought tolerance (r > 0.82, p < 0.01). It is interesting to note that, even in the experimental settings as reported by us earlier, the native biofilms on control plants comprised around 6% D-GA (Table 2) (Timmusk, Copolovici et al. 2019). EPS biosynthesis involves a very sophisticated network dependent on complex factors (Schmid, Sieber et al. 2015). Uronic acid moieties affect the physiological and biological abilities of polysaccharides, e.g. the uronates with high content of EPS, such as alginate and xanthan, result in increased water holding capacity (Liang and Wang 2015). Uronic acid backbones lead to changes in other sugar backbones, which eventually results in alteration of their properties and bioactivity (Xu, Bi et al. 2014).

Compare to proteins, the biosynthesis of EPS is not template driven, and polysaccharides occur as heterogeneous mixtures of high complexity. MALDI TOF has been developed as a successful tool for the analysis of biopolymers and is a valuable tool for comparison of polysaccharide profiles (Kailemia, Ruhaak et al. 2014, Lopez-Garcia, Garcia et al. 2016). The ions produced are mostly single charges molecular ions and the mass spectra can be used for distribution analysis of polysaccharides. Aqueous extracts can be successfully used without further purification (Kailemia, Ruhaak et al. 2014, Lopez-Garcia, Garcia et al. 2016) A26Sfp culture filtrate EPS production was compared to the SN induced strain EPS production. The results confirms the EPS titre results (Table 2). While the SN induced A26Sfp low weight EPS chains show relatively small increase the higher weight EPS chain production increases more than ten times (Fig 2, Table 3).

Silica particle effects, on plant and bacterial performance under various stress conditions, have been studied earlier and as one of the effects bacterial count/ biomass increase is reported (Karunakaran, Suriyaprabha et al. 2013, Rangaraj, Gopalu et al. 2014, Luyckx, Hausman et al. 2017). Our results show that while SNs improved the EPS production of both harsh environment strains used, the SN treatments didn’t increase bacterial count (Table 2). The result suggests diverse mechanism involvement and indicates that harsh environments isolates may have advantage of their enhanced EPS production. Microorganisms in their native environments are subjected to various fluctuations in environmental conditions. It has been suggested that the ecological ‘success’ of EPS depends on their potential to beneficially influence the bacterial adaptation to the environment (Geoghegan, Andrews et al. 2008). The EPS matrix serves as the microbial interface with the environment (Costa, Raaijmakers et al. 2018). EPS molecules with high WHC (Liang and Wang 2015, Costa, Raaijmakers et al. 2018)can mechanically protect against water stress and were suggested by us as a mechanism of the drought tolerance enhancement of harsh environment rhizobacterial strains (Nevo 2012, Timmusk, Kim et al. 2015). Our results show that SN presence leads to EPS synthesis with higher D-GA content. The EPS protective layer with high WHC on the root surface could re-establish water potential gradients under drought stress (Table 3). Even though it is a most probable mechanism earlier reported by several groups (Timmusk and Behers 2019, Timmusk, Copolovici et al. 2019, Timmusk and Zucca 2019) it is likely not to be the only one, especially in view of the extreme habituation in the rhizosphere of wild progenitors of cereals. Production and accumulation of osmoregulatory compounds and maintenance of low internal pressure via efflux pumps are the additional potential mechanisms, all regulated by complex gene expression machinery. While it is known that the internal osmotic pressure in biofilms is primarily generated by the EPS, the possibility of a macromolecular osmolyte secreted and maintained by the cells, in response to the influence of osmotic pressure gradients on the growth characteristics of biofilms, remains largely underexplored. It is generally believed that reactions involved in gene translation are ionic-dependent and can be explained in electrostatic terms (Douzou 1994). Studies indicate that there is a gap between the rough picture of the mechanism of ionic regulation and detailed behaviour of reactions at the molecular level (Douzou 1994, Morbach and Kramer 2002, Zhang and al 2013, Falghoush, Beyenal et al. 2017, Srinivasan and al 2018). While the complete signal transduction pathways of the cells’ response to osmotic challenge is far from being understood, the sequence of events starts with a signal from an osmosensory receptor. The signal is then transduced to transport systems leading to the response of the cell resulting in osmoresponsive changes is gene expression (Timmusk, Kim et al. 2015).

Formerly we have shown that nanoparticles stabilise plant growth promoting rhizobacterial attachment and colonization(Timmusk, Seisenbaeva et al. 2018). This result is translated into plant biomass accumulation and growth improvements in stressful environments. The present work has revealed that SNs significantly increase EPS formation. The physical property of the matrix, EPS content, increases, which in turn increases WHC and osmotic pressure. Increased WHC and internal hyperosmolarity are likely to regulate the behaviour of the cells. The microscope studies show that in the presence of SN particles, bacterial vitality and morphology are changed (Fig 4 and Video S1). This indicates that SNs are associated with significant reprogramming in bacteria, and the treatment initiates novel physiological properties of the bacteria that are not present in cells without SN treatment.

It is clear that understanding mechanisms of the desiccation tolerance induced by the harsh environment isolate A26 and its mutant A26Sfp may not follow from the study of the expression of a few genes or even that of the total genome. To grasp the basis of the complete system will require detailed functional genomic, proteomic and physiological and physical studies in the framework of silica particles. It has been often suggested that the whole can’t be described as a summa of particles and holobiotic system reductionist approach has been a topic of controversy(Anderson 1972). A comprehensive analysis of regulatory genes and transcription factors based on the transcriptome of isolates exposed to osmotic stress treatments is needed. Across this range of work, high resolution microscopy of the structure of the bacterial surface complex of biomolecules and cellular organelles, should be applied in order to gain insight into how stress-tolerance is coupled with the morphology of the bacteria

## CONCLUSIONS

Our results with the commercial SNs agree with former results with natural soil silicas and conform that the plant growth-promoting rhizobacteria beneficial effects are induced by SN treatment. The findings encourage formulation of cells considering microencapsulation with silica containing materials for field applications. Future research has to be performed on magnitude, stability, dynamics and functions of silica pools in agricultural soils.

## Supporting information

A26Sfp midlogaritmic culture with SN

A26Sfp midlogaritmic culture without SN

## ACKNOWLEDGEMENTS

The work was performed with financial support from the Carl Tryggers Stiftelse för Vetenskaplig Forskning, Swedish Research Council 2017-5224, FORMAS 222-2014-1326 and SIDA Biotech (to ST).

## SUPPLEMENTARY MATERIAL

Video S1

### Video recordings of mid logarithmic bacterial culture grown with and without of SN particles

Typical performance of A26 and A26Sfp bacteria grown with (A) and without of SN particles. Recording for A26Sfp is shown See Material and Methods.

### METHODS

#### Bacterial growth and culture conditions

*P*. *polymyxa* A26^4^ originates from wild barley rhizosphere from Facing Slope at the natural laboratory called the Evolution Canyon (Table 1). The *P*. *polymyxa* A26 Sfp-type 4-phosphopantetheinyl transferase deletion mutant strain A26Sfp (A26△sfp), was generated as previously described^9,33,34,36^. Stock cultures were stored at −80 °C and streaked for purity on half strength tryptic soy agar (1/2 TSA). All bacterial strains were grown until mid logarithmic growth phase in half-strength tryptic soy broth (1/2 TSB; pH 6.2) at 30 ±2 °C.

Cultures were centrifuged and pellets re-suspended in PBS as described earlier (Abd El-Daim, Haggblom et al. 2015). Finally, the cultures were adjusted to 10^6^ cells /ml.

#### Bacterial growth in the presence of nanoparticles

Commercial Ludox^®^ TM50 Sigma Aldrich particles were used in the study (Various 2006, Orts-Gil, Natte et al. 2011). Strains A26, A26Sfp were grown in 1/2 TSB with 50 μg ml^-1^ particles. The SNs were deagglomerated by ultrasound for 1-5 min and mixed with growth medium prior to bacterial inoculation. Culture density was determined by colony forming unit analysis (CFU). For the mock treatment, 1/2 TSB with 50 μg ml^-1^ SNs was used.

#### EPS and D-GA content evaluation

EPS were isolated from the A26, A26SN and A26Sfp, A26SfpSN cultures and root biofilms. The bacterial cultures and root biofilms were prepared as described above. The root biofilm bacteria were re-isolated by selective plating, confirmed by PCR as described earlier (Timmusk, Abd El-Daim et al. 2014, Timmusk, Kim et al. 2015) and quantified. EPS extraction was performed as described earlier (Timmusk, Copolovici et al. 2019) (Rutering, Schmid et al. 2016). Briefly, the biofilms were harvested washing by PBS. Bacterial cultures were diluted 1:5 with distilled water and centrifuged for 30 min at 17,600 g at 20 °C to separate cells. Then, EPS were extracted twice by precipitation, slowly pouring the supernatant into two volumes of isopropanol while stirring at 200 rpm. The filtered polysaccharide was suspended in a digestion solution consisting of 0.1 M MgCl2, 0.1 mg/ml DNase and 0.1 mg/ ml RNase solution, and incubated for 4 h at 37 °C. Samples were extracted twice with phenol-chloroform, lyophilised using a Virtis SP Scientific 2.0 freeze dryer, taken to the initial volume and dialysed against distilled water. Analysis of uronic acid content was performed as described earlier by Mojica et al 2007 (Mojica, Elsey et al. 2007) with small modifications. Briefly, the EPS pellets were weighed and dissolved in 200 μl of deionised water. Potassium sulfamic acid (4M, pH 1.6) was added to the EPS solution and mixed by vortexing. Then sodium tetraborate solution in concentrated sulfuric acid (0.0125M) was added. The solutions were incubated for 5 min in a 100 °C water bath, cooled on ice for 3 min and centrifuged at 2,000 rpm for 10 min, after which 20 μl hydroxyophenol solution (0.15% v/v) was added to the supernatant. The solution was then mixed gently and the absorbance read at 520 nm. Each data point represents the average of twenty replicate measurements.

#### MALDI mass spectrometry

Angiotensin II (1045.5 u), ACTH clip 18–39 (2464.2 u), bovine insulin (5733.5 u) and equine cytochrome c (12360.1 u) were used for external calibration of the mass spectrometer. All calibrants (1–5 pmol/μl) were dissolved in 0.1% trifluoroacetic acid (TFA) in ultrapure Milli-Q Plus water. 2,5-dihydroxy benzoic acid (DHB) was used as MALDI matrix. The matrix was dissolved in HPLC-grade acetronitrile (1 g/l for the seed layer) or in 0.1% TFA in acetonitrile/ultrapure water [1:1 v/v] (10 g/l, saturated for the sample/matrix mixture).

All samples were prepared with the seed layer method. First, a matrix seed layer was created by depositing a droplet (1 μl) of a 1 g/l solution of matrix dissolved in acetonitrile on a highly polished, stainless steel sample probe. Thereafter, the 10 g/l matrix and sample solutions were mixed in a test tube 1:1, and a droplet (1 μl) of sample/matrix was deposited on the matrix seed layer. Samples were then left to dry totally in air.

All MALDI analyses were performed with an upgraded Reflex II MALDI-TOF mass spectrometer (Bruker-Franzen Analytic Gmbh, Bremen, Germany). Samples were irradiated with a 337 nm nitrogen and a laserspot ~10–20 μm in diameter. All spectra were acquired in the reflectron mode at an accelerating voltage of 20 kV. Mass spectra were analyzed using the software provided by Bruker (Niedermeyer and Strohalm 2012). All spectra shown were calibrated using external calibration with a mass deviation of within 0.08%.

#### Light Microscopy

Bacterial strains were grown until mid-logarithmic phase of growth with and without SNs as described above. A Celestron PentaView LCD Digital Microscope was used to collect images and record videos of the bacterial growth in logarithmic phase.

#### Cryo-Electron Microscopy

A26 and A26Sfp strains were grown in half strength TSB in the presence and absence of SNs as described above. At the mid-logarithmic phase of growth, the cultures were sieved in order to collect the bacterial clumps using PELCO prep-eze individual wells (Ted Pella). The trapped bacterial clumps were then washed twice in PBS, sonicated and adjusted to 10^3^ cells per ml. For comparison, from the cultures that did not form clumps, the bacterial pellets were washed twice in PBS and diluted to 10^3^ per ml.

3 μl of bacteria solution at a concentration of 10^3^ per ml was applied to glow-discharged (40 s, 20mA using Pelco easyGlow) C-Flat 200 mesh R2/2 (thick) grids. Using a Vitrobot Mark IV (Thermo Fisher Scientific) operated at 4°C and 100% humidity, grids were blotted for 4 s before being flash-frozen in liquid ethane.

All TEM data were collected with a Talos Arctica (Thermo Fischer Scientific) microscope operated at 200kV and a Falcon III camera (Thermo Fisher Scientific) operated in integrating mode. Images were collected at 8700x nominal magnification using approximately 2 electrons/A2 total dose and a defocus of – 30 μm.

#### Osmolarity assay

Strains A26, A26Sfp were grown until mid-logarithmic growth phase in half-strength tryptic soy broth as described above and the cultures were adjusted to 10^6^ cells /ml. The osmotic pressure of the solution was measured with an Advanced Instruments Freezing Point Osmometer (model 3250). The contribution of the polymer to the osmotic pressure was calculated by subtracting the measured osmotic pressure of the pure medium.

#### Evaluation of A26 and A26Sfp EPS water holding capacity (WHC)

Bacterial cultures were grown and harvested, and polysaccharides isolated, as described above. 600 mg of A26 and A26Sfp EPS, were mixed with 100 g peat soil (Sol mull, Hasselfors) and determined at different wetting cycles for 24 h. The soil biopolymer mixture was then allowed to drain for 30 min and the weight after saturation was recorded. Following the wet weight estimation the mixture was dried in an oven, cooled in a desiccator and weighed again. Each experiment was carried out in triplicate. WHC was calculated as percentage of water holding capacity: = gain in weight at saturation point/dry weight of the soil x 100.

#### Plant treatment

Winter wheat (Stava) seeds were surface sterilised by a 60 s wash in 99% ethanol, followed by a 6 min wash in 3% sodium hypochlorite solution, and a wash in 99% ethanol, and rinsed several times with sterilized water. For germination the sterilised seeds were placed in Petri dishes with one sheet of filter paper moistened with 5 mL of distilled water. The dishes were kept in a dark incubator at 20 °C until germination. A germination paper (4.5 × 3.0 cm) was moistened with 10 mL of sterile water.

Uniformly germinated wheat seeds were grown in pots filled with peat soil (Sol mull, Hasselfors) and incubated in an MLR-351H (Phanasonic, IL, USA) growth chamber at 24/16 °C (day/night) temperature, and 16 h photoperiod at 250 μmol m^-2^ s^-1^ (Table 1). On the fifth day of seedling growth, the PGPR inoculation was performed by adding 1 ml of bacteria at 10^6^ cells /ml grown as described above. The soil moisture was adjusted to 75% of water holding capacity. Soil moisture (12.5% of soil dry weight) was kept constant during the first 8 days of seedling growth. Soil volumetric water content was evaluated using 5TE soil moisture sensors (Decagon Devices, Inc).

The other set of plants was grown hydroponically as described earlier(Zheng, Wen et al. 2019). Briefly, uniformly germinated seeds were placed in plastic boxes (10 × 5 × 2.5 cm containing 1.5 L of nutrient solution, pH = 5.8) and supported vertically. The nutrient solution was refreshed every third day. On the fifth day of seedling growth, the PGPR inoculation was performed by adding 1 ml of bacteria 10^6^ cells /ml grown as described above (Table 1).

After 8 days of growth ten seedlings were carefully removed from the growth environment (peat and hydrophonic culture) and root homogenization and bacterial identification and quantification were performed. In addition to bacterial population assay, EPS and D-GA content was evaluated from the seedlings grown in hydroponic culture (Table 1).

The plants grown in soil were exposed to seven-day-long drought stress after 8 days of growth. The experiment was performed in four replicates, each consisting of 6 plants. Plant biomass is expressed as shoot dry mass. Wheat shoot samples were dried at 105 °C to a constant mass, cooled and weighed.

#### Data confirmation and validation

To ensure reproducibility, three biological replicates of every single PGPR treatment were performed, if not stated otherwise. Twenty biological replicates of each D-glucuronate detection experiment were performed. Replicated data were studied for normal distribution and analysed by Unscrambler X15.1 and MiniTab17. SN treatment effects were considered statistically significant at *p* ≤ 0.05. Linear regressions (Unscrambler X15.1) were used to determine the relationships between seedling biomass accumulation and D-glucuronic acid content.

**Table S1.**
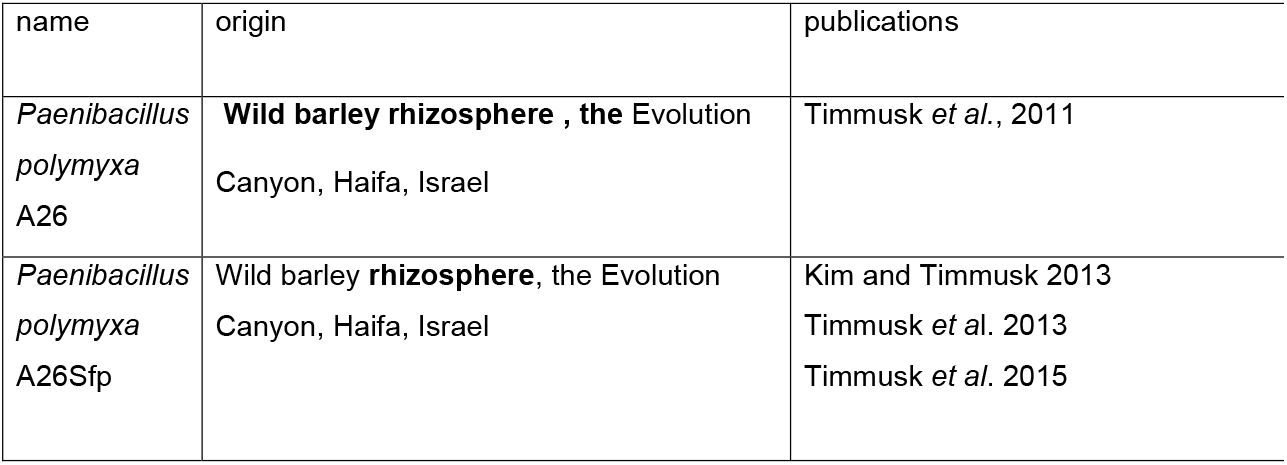
Strains used in the study

| name | origin | publications |
| --- | --- | --- |
| *Paenibacillus polymyxa* A26 | **Wild barley rhizosphere , the** Evolution Canyon, Haifa, Israel | Timmusk *et al.*, 2011 |
| *Paenibacillus polymyxa* A26Sfp | Wild barley **rhizosphere**, the Evolution Canyon, Haifa, Israel | Kim and Timmusk 2013<br>Timmusk *et al.* 2013<br>Timmusk *et al.* 2015 |

## Notes

### Competing Interest Statement

The authors have declared no competing interest.

## REFERENCES

Abd El-Daim, I., P. Haggblom, M. Karlsson, E. Stenstrom and S. Timmusk (2015). “*Paenibacillus polymyxa* A26 Sfp-type PPTase inactivation limits bacterial antagonism against *Fusarium graminearum* but not of *F. culmorum*” Front. Plant Sci.: 1–8.

Anderson, P. W. (1972). “More is different” Science vol 177.

Bandeppa, S., S. Paul, J. K. Thakur, N. Chandrashekar, D. K. Umesh, C. Aggarwal and A. D. Asha (2019). “Antioxidant, physiological and biochemical responses of drought susceptible and drought tolerant mustard (Brassica juncea L) genotypes to rhizobacterial inoculation under water deficit stress.” Plant Physiol Biochem 143: 19–28.

Costa, O. Y. A., J. M. Raaijmakers and E. E. Kuramae (2018). “Microbial extracellular polymeric substances: ecological function and impact on soil aggregation.” Front Microbiol 9: 1636.

Dai, A. (2012). “Increasing drought under global warming in observations and models.” Nat Clim Change 3: 52–58.

Douzou, P. (1994). “Osmotic regulation of gene action.” Proc Natl Acad Sci U S A 91(5): 1657–1661.

Falghoush, A., H. Beyenal, T. Besser, A. Osmland and D. Call (2017). “Osmotic componds enhance antibiotic efficacy against *Acinetobacter baumannii* biofilm communities” Applied and Environmental Microbiology.

Fitzpatrick, C. R., J. Copeland, P. W. Wang, D. S. Guttman, P. M. Kotanen and M. T. J. Johnson (2018). “Assembly and ecological function of the root microbiome across angiosperm plant species.” Proc Natl Acad Sci U S A 115(6): E1157–E1165.

Fitzpatrick, C. R., Z. Mustafa and J. Viliunas (2019). “Soil microbes alter plant fitness under competition and drought.” J Evol Biol 32(5): 438–450.

Geoghegan, M., J. S. Andrews, C. A. Biggs, K. E. Eboigbodin, D. R. Elliott, S. Rolfe, J. Scholes, J. J. Ojeda, M. E. Romero-Gonzalez, R. G. Edyvean, L. Swanson, R. Rutkaite, R. Fernando, Y. Pen, Z. Zhang and S. A. Banwart (2008). “The polymer physics and chemistry of microbial cell attachment and adhesion.” Faraday Discuss 139: 85–103; discussion 105-128, 419–120.

Gilbert, S. F. (2016). “Developmental Plasticity and Developmental Symbiosis: The Return of Eco-Devo.” Curr Top Dev Biol 116: 415–433.

Gilbert, S. F. (2019). “Developmental symbiosis facilitates the multiple origins of herbivory.” Evol Dev: e12291.

Gilbert, S. F., T. C. Bosch and C. Ledon-Rettig (2015). “Eco-Evo-Devo: developmental symbiosis and developmental plasticity as evolutionary agents.” Nat Rev Genet 16(10): 611–622.

Hristozov, D., L. Pizzol, G. Basei, A. Zabeo, A. Mackevica, S. F. Hansen, I. Gosens, F. R. Cassee, W. de Jong, A. J. Koivisto, N. Neubauer, A. Sanchez Jimenez, E. Semenzin, V. Subramanian, W. Fransman, K. A. Jensen, W. Wohlleben, V. Stone and A. Marcomini (2018). “Quantitative human health risk assessment along the lifecycle of nano-scale copper-based wood preservatives.” Nanotoxicology 12(7): 747–765.

Kailemia, M. J., L. R. Ruhaak, C. B. Lebrilla and I. J. Amster (2014). “Oligosaccharide analysis by mass spectrometry: a review of recent developments.” Anal Chem 86(1): 196–212.

Karunakaran, G., R. Suriyaprabha, P. Manivasakan, R. Yuvakkumar, V. Rajendran, P. Prabu and N. Kannan (2013). “Effect of nanosilica and silicon sources on plant growth promoting rhizobacteria, soil nutrients and maize seed germination.” IET Nanobiotechnol 7(3): 70–77.

Liang, T. W. and S. L. Wang (2015). “Recent advances in exopolysaccharides from *Paenibacillus spp.:* production, isolation, structure, and bioactivities.” Mar Drugs 13(4): 1847–1863.

Liepe, K. J., A. Hamann, P. Smets, C. R. Fitzpatrick and S. N. Aitken (2016). “Adaptation of lodgepole pine and interior spruce to climate: implications for reforestation in a warming world.” Evol Appl 9(2): 409–419.

Lopez-Garcia, M., M. S. Garcia, J. M. Vilarino and M. V. Rodriguez (2016). “MALDI-TOF to compare polysaccharide profiles from commercial health supplements of different mushroom species.” Food Chem 199: 597–604.

Luyckx, M., J. F. Hausman, S. Lutts and G. Guerriero (2017). “Silicon and Plants: Current Knowledge and Technological Perspectives.” Front Plant Sci 8: 411.

Mojica, K., D. Elsey and M. J. Cooney (2007). “Quantitative analysis of biofilm EPS uronic acid content.” J Microbiol Methods 71(1): 61–65.

Morbach, S. and R. Kramer (2002). “Body shaping under water stress: osmosensing and osmoregulation of solute transport in bacteria.” ChemBioChem.

Naylor, D. and D. Coleman-Derr (2017). “Drought Stress and Root-Associated Bacterial Communities.” Front Plant Sci 8: 2223.

Nevo, E. (2012). ““Evolution Canyon,” a potential microscale monitor of global warming across life.” Proc Natl Acad Sci U S A 109(8): 2960–2965.

Niedermeyer, T. H. and M. Strohalm (2012). “mMass as a software tool for the annotation of cyclic peptide tandem mass spectra.” PLoS One 7(9): e44913.

Orts-Gil, G., K. Natte, D. Drescher, H. Bresch, A. Mantion, J. Kneipp and W. Osterle (2011). “Characterization of silica nanoparticles prior to in vitro studies.” J. Nanopart Res.

Rangaraj, S., K. Gopalu, Y. Rathinam, P. Periasamy, R. Venkatachalam and K. Narayanasamy (2014). “Effect of silica nanoparticles on microbial biomass and silica availability in maize rhizosphere.” Biotechnol Appl Biochem 61(6): 668–675.

Rice, R., A. Kallonen, J. Cebra-Thomas and S. F. Gilbert (2016). “Development of the turtle plastron, the order-defining skeletal structure.” Proc Natl Acad Sci U S A 113(19): 5317–5322.

Robinson, C. D., H. S. Klein, K. D. Murphy, R. Parthasarathy, K. Guillemin and B. J. M. Bohannan (2018). “Experimental bacterial adaptation to the zebrafish gut reveals a primary role for immigration.” PLoS Biol 16(12): e2006893.

Rutering, M., J. Schmid, B. Ruhmann, M. Schilling and V. Sieber (2016). “Controlled production of polysaccharides-exploiting nutrient supply for levan and heteropolysaccharide formation in Paenibacillus sp.” Carbohydr Polym 148: 326–334.

Schmid, J., V. Sieber and B. Rehm (2015). “Bacterial exopolysaccharides: biosynthesis pathways and engineering strategies.” Front Microbiol 6: 496.

Shimaila, A. and B. R. Glick (2019). Plant-bacterial interactions in management of plant growth under abiotic stress. New and future developments in microbial biotechnology and bioendineering. B. R. Glick. Elsevier B V.

Srinivasan, S. and e. al (2018). “atrix production and sporulation in Bacilllus subtilis biofilms localize to propagating wave fronts.” Biophysical Journal.

Sukweenadhi, J. and e. al (2015). “Paenibacillus yonginensis DCY84T induces changes in Arabidopsis thaliana gene expression.” Microb Res.

Timmusk, S., I. A. Abd El-Daim, L. Copolovici, T. Tanilas, A. Kannaste, L. Behers, E. Nevo, G. Seisenbaeva, E. Stenstrom and U. Niinemets (2014). “Drought-tolerance of wheat improved by rhizosphere bacteria from harsh environments: enhanced biomass production and reduced emissions of stress volatiles.” PLoS One 9(5): e96086.

Timmusk, S. and L. Behers (2019). “Nanotechnology, is it something useful for future agriculture?” Atlas of Science http://atlasodscience.org.

Timmusk, S., D. Copolovici, L. Copolovici, T. Teder, E. Nevo and L. Behers (2019). “*Paenibacillus polymyxa* biofilm polysaccharides antagonise *Fusarium graminearum* DOI:10.1038/s41598-018-37718-w” Nature Sci. Rep..

Timmusk, S., I. El Daim, L. Copolovici, A. Kannaste, L. Behers, E. Nevo, S. G. E. Stenstrom and U. Niinemets (2014). “Drought-tolerance of wheat improved by rhizosphere bacteria from harsh environments: enhanced biomass production and reduced emissions of stress volatiles.” PloS ONE 113

Timmusk, S., S. B. Kim, E. Nevo, I. Abd El Daim, B. Ek, J. Bergquist and L. Behers (2015). “Sfp-type PPTase inactivation promotes bacterial biofilm formation and ability to enhance wheat drought tolerance.” Front Microbiol 6: 387.

Timmusk, S., V. Paalme, T. Pavlicek, J. Bergquist, A. Vangala, T. Danilas and E. Nevo (2011). “Bacterial distribution in the rhizosphere of wild barley under contrasting microclimates.” PLoS One 6(3): 1–8.

Timmusk, S., G. A. Seisenbaeva and L. Behers (2018). “Titania (TiO2) nanoparticles enhance the performance of growth-promoting rhizobacteria DOI: 10.1038/s41598-017-18939-x.” Nature Sci Rep.

Timmusk, S. and E. G. Wagner (1999). “The plant-growth-promoting rhizobacterium *Paenibacillus polymyxa* induces changes in *Arabidopsis thaliana* gene expression: a possible connection between biotic and abiotic stress responses.” Mol Plant Microbe Interact 12(11): 951–959.

Timmusk, S. and C. Zucca (2019). “The plant microbiome as a resource to increase crop productivity and soil resilience: A systems approach.” Journal of Cameroon Academy of Sciences Vol 14, No 3 DOI: 10.4314/jcas.v14i3.2.

Various (2006). “Synthetic amorphous silica. European centre for ecotoxicology and toxicology for chemicals.” ECETOC JACC No 51.

Vega, N. M. (2019). “Experimental evolution reveals microbial traits for association with the host gut.” PLoS Biol 17(2): e3000129.

Xu, L., D. Naylor, Z. Dong, T. Simmons, G. Pierroz, K. K. Hixson, Y. M. Kim, E. M. Zink, K. M. Engbrecht, Y. Wang, C. Gao, S. DeGraaf, M. A. Madera, J. A. Sievert, J. Hollingsworth, D. Birdseye, H. V. Scheller, R. Hutmacher, J. Dahlberg, C. Jansson, J. W. Taylor, P. G. Lemaux and D. Coleman-Derr (2018). “Drought delays development of the sorghum root microbiome and enriches for monoderm bacteria.” Proc Natl Acad Sci U S A.

Xu, X., D. Bi, X. Wu, Q. Wang, G. Wei, L. Chi, Z. Jiang, T. Oda and M. Wan (2014). “Unsaturated guluronate oligosaccharide enhances the antibacterial activities of macrophages.” FASEB J 28(6): 2645–2654.

Zhang, W. and e. al (2013). “Nutrient depletion in *Bacillus subtilis* biofilms triggers matrix production.” New Journal of Physics.

Zheng, X., X. Wen, L. Qiao, J. Zhao, X. Zhang, X. Li, S. Zhang, Z. Yang, Z. Chang, J. Chen and J. Zheng (2019). “A novel QTL QTrl.saw-2D.2 associated with the total root length identified by linkage and association analyses in wheat (Triticum aestivum L.).” Planta 250(1): 129–143.

